# Protocol for clustering of non-unified protein sequences through memory-map guided deep learning

**DOI:** 10.1101/2020.08.15.252114

**Authors:** Om Prakash

## Abstract

Protocol established and validated for clustering of non-unified protein sequences through memory-map guided deep learning. Data evaluated belongs to the disease causing proteins/genes from human hormonal system. Possibilities for future experiments validation was found for genes as: ACTHR, AGMX1, ATK, BPK, DPDE3, ERBA2, FSHB, GH1, GHSR, GNAS1, GSP, HANF, LCGR, LGR2, LGR3, LHRHR, NR1A2, PKR1, PRKAR1, RNF216, SBP2, SECISBP2, THR1, THRB, TPIT, TRIAD3, TSE1, UBCE7IP1, XAP2, and ZIN. Protocol is recommended for implementation with small to large dataset (protein/ DNA/ RNA sequences of unified or non-unified length) with unclassified data flags.

## INTRODUCTION

Order of amino acid sequence is the basis of 3D structure and functional specificity of protein. 3D structure of protein sequence represents responsible structure for function of protein. Protein sequences are known to be classified on the basis of sequences of same stretch of length through multiple sequence alignment. But in today’s scenario, when next-generation-sequencing data are being provided into a large scale, then sequences of unclassified data with non-unified length are becoming a challenge. Unclassified as well as non-unified length of protein sequences can’t be directly used for clustering or classification which can be performed on the basis of their functions. To handle such complication, sequence memory map can be a suitable solution for pre-processing of non-unified sequence data. Sequence memory map has been already defined to capture specific order of existence of amino acids or nucleotide into sequence. Therefore it can be used as an important method to map the sequences of non-uniform length. Since the mapping algorithm conserves the relative location of string elements into memory map, therefore it becomes a representative of structure-to-function link between genotype and phenotype. Details about algorithm can be accessed from *Prakash, 2020* ‘doi: https://doi.org/10.1101/2020.03.05.979781’. Deep learning involves learning of artificial neural network with pre-processed data with customized handling of biological essence of functionality. Here deeplearning theme was implemented to capture the theme of linking genotype-to-phenotype for clustering of unclassified data.

The study has been found to be covered through supported references as: As it is well known that though BLAST (Basic Local Alignment Search Tool) the query sequence is clustered with sequences from databases. Here sequence matching score is defined on the basis of scoring matrices (*Altschul et.al. 1990*). By using memory map, the customized dataset of sequences can be scored in un-supervised manner. This can be a representation of alignment of memory maps. Such map alignments are feasible through deep-learning. Deep clustering has been priory implemented for clustering of large protein sequence sets. Therefore deep-learning based scoring of memory map can be an efficient method (*Hauser et. al. 2016*). At large scale, protein sequences have been classified in protein families. In continuation of similar themes, unclassified non-unified protein sequences can be further processed for network mapping (*Finn et. al. 2016*). Sequence memory maps can also be used for comparison of mutational variations (*Hopf et. al. 2017*). Clustering algorithms are being advanced for handling of large as well as complex datasets (*Fu et. al. 2012*). Fast clustering algorithms are available for metagenomic sequence analysis (*Li et. al. 2012*).

Therefore in present work, a protocol has been presented for clustering of non-unified protein sequences through memory-map guided deep learning.

## MATERIALS & METHODS

### Dataset for mapping

Protein set was collected from UniProt, a universal protein resource. Dataset included known protein sequences (and respective UniProt IDs) from hormone signaling systems involved in multiple diseases of human. The set included 33 unique protein objects.

### Protocol for Deep-Learning

Overall workflow of protocol has been shown in figure 1; which includes (i) preparation of sequence memory map; (ii) flattening of memory map; (iii) normalization of data; (iv) bipolar rotational supervised learning of ANN models; and (v) post-analysis of model outputs for network mapping.

**Figure 1.**
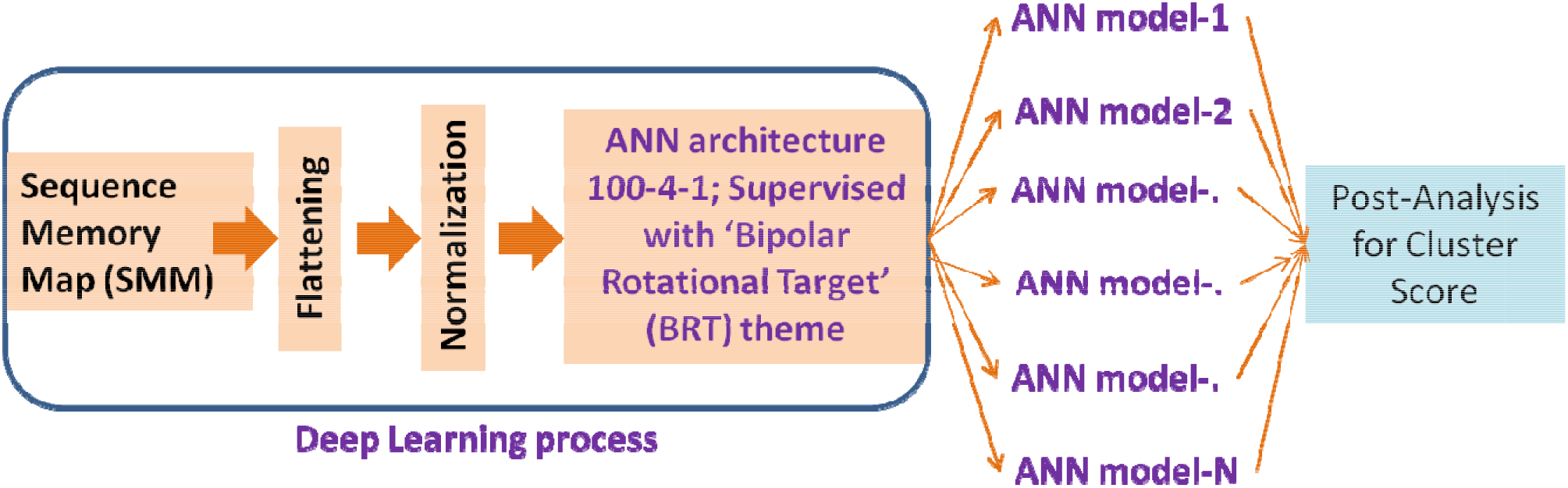
Workflow of deep learning with post processing for clustering of dataset

#### Step1: Development of Sequence Memory Map (SMM)

Initial requirement for starting the algorithm was a set of bio-strings (here protein sequence), each string out of set was transformed into ‘sequence memory map’. Firstly SMM was initialized as zero matrix of order u x w. where u is the number of unique characters in the data type (here for protein sequence u = 20). Number of columns w represented window-length used for map enrichment. Zero matrix was enriched with elemental observation of string. Each element of string throws a value into matrix in respect of their position in string as well as unique character. Memory matrix has been filled with respective value of string element. Details of memory mapping are as following (‘doi: https://doi.org/10.1101/2020.03.05.979781’): Let O_uxv_ be a SMM zero matrix of order u x v. Let S = {s_n_} be a string of length n(s). Let an ordered set, where u1,u2,u3,u4,u5,u6,u7,u8,u9,u10,u11,u12,u13,u14,u15,u16,u17,u18,u19,u20 are denoted by A,C,D,E,F,G,H,I,K,L,M,N,P,Q,R,S,T,V,W,Y respectively. Here w denoted window size. Let the smallest integer ≥. Let W_nc_ denote the rearranged string S in the matrix of order d x c (*Blank element will be considered, if required during rearrangement*).

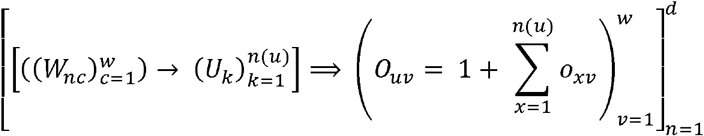

#### Step-2: Flattening of SMM

Memory map was flattened to vector form. Set of flattened SMM was normalized; which was further used as inputs of ANN model.

#### Step-3: Customized training of Artificial Neural Network

ANN architecture of 100-4-1 was implemented for model development. Although ANN architecture was supervised; but customized training was processed with ‘bipolar rotational target’ (BRT) theme. BRT was introduced as that each sample was trained with both binary values ‘0’ and ‘1’, but in rotational manner for multiple times. Multiple models were prepared with multiple seed values.

#### Step-4: Post-training processing

Multi-model outputs were processed to calculate cluster-score for each sample. Cluster-score is a z-score values identified through nested processing of Multi-model outputs. Cluster score has been defined as:

Let’s Sample set: {S_1 to M_}_N_

Where ‘N’ is number of samples (S); and each sample has ‘M’ number of outputs from ‘M’ ANN-models. For each sample vector; mean {μ_N_} and standard deviation {σ_N_} were calculated. Furthermore mean (μ’) and standard deviation (o’) was calculated for {(μ/σ)_N_}

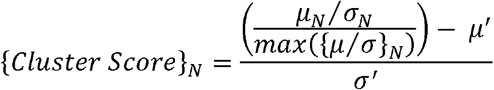

#### Step-5: Classification of samples in two classes

On the basis of Cluster-score

#### Step-6: Network Graph with node values as cluster-score

Cluster-score used as target tags for network mapping. Since ANN model capture core relation between data points, therefore it was supposed to get the highest linked (i.e. experimentally proved) interactions into zone of least distant nodes; and low linked interactions in zone of maximal distant nodes. This assumption was cross validated with experimentally with known evidences.

## RESULTS & DISCUSSION

Overall, a theoretical mapping of biostrings was used for representation of their expression, which was explained for bridging genotype to phenotype. Main algorithm was tested with the protein data set. This algorithm receives input as a set of biostrings, as protein or nucleotide sequence. SMM algorithm develops a memory map on the basis of sequences provided as input. Other details can be found from the referenced literature (‘doi: https://doi.org/10.1101/2020.03.05.979781’). SMM algorithm was basically used for converting non-unified sequences into unified memory maps. Each map has elemental component of integer values, and whole map can be represented as matrix.

For deep-learning, preparation of unified matrix was the first part of deep learning, before the implementation of artificial neural network. Second step was to convert the unified metrics into the inputs of artificial neural network. For which, each matrix was flattened into vector. Accordingly each matrix was flattened. At this stage, each vector element contained integer values. In this way by processing each of the protein, a set of vectors of unified length was obtained. Next step was to normalize the data prepared till now. Here global normalization was implemented. For global normalization, whole data set was normalized with the single maximum value. Here each element of the vector was considered as input for nodes at artificial neural network. Next stage was to implement artificial neural network. Since the data was unclassified that’s why there was no class value. In this study our motivation was to categories the unclassified data. Therefore a logical intervention was implemented with artificial neural network. Although network architecture was defined in its traditional view but the target value for supervision of the model was defined with a logical rules. Since our aim was to classify the data, therefore initially it was assumed that each string belongs to the binary classes as ‘0’ and ‘1’. Therefore, two binary flags were linked with each sample, but it was included into a circular replacement during the multiple training of the model. In this way each sample was participated in the training of artificial neural network with both supervision targets ‘0’ and ‘1’. Multiple ANN training models were prepared with circular network output, and training outputs were recorded.

In post-training process, outputs from multiple trained artificial neural network models were analyzed to classify the data. Since each sample was observed at the mean value of the two targets ‘0’ and ‘1’. Therefore a customized cluster scoring parameter was defined on the basis of mean and standard deviation of the outputs from multiple models. Cluster score classifed the data into two parts. These results include two major interpretations: First that non-classified data was categorized into two sections; secondly each data point was flagged with floating value from multiple trained artificial neural networks, which can be further used for training of the system of multiple samples. The floating point values for each protein was used to prepare network graph for possible interactions among the proteins considered in the hormones related genes for diseases. For network graph preparation, with Networkx python package, distance between the each node was used as edge weight. Edge weight was used as normalized value between 0 and 1. Furthermore, network graph was filtered to identify the set of genes with similar or differential performance. To validate the classified data found from filtering the network graph, each sample was evaluated with the information available in literature through STRING database.

Total 33 proteins sequences (**Supplementary Table 1**) were considered from UniProt database, regarding hormonal proteins involve in various diseases. These proteins were belonging to the genes related with hormones in human system. Since no supervision was present to classify such unclassified non-unified sequences.

### Cross-validation of clustering based on experimental evidences

Two types of network graphs were plotted. First was, STRING database based network graph, plotted with 33 protein sequences. Second was, network graph plotted on the basis of node values adopted from customized deep-learning. Since first network graph showed experimentally proved gene interactions, along with other methods as text mining, co-expression, protein homology etc. (Figure 2). Therefore experimentally approved interactions were considered as reference for cross validation of efficiency of protocol adopted for deep-learning. Figure 3 showed the network’s circular graph with all combinations including edge weight of 0 to 1. Here node values were calculated through deep-learning based multiple models. Inter-node edge weight was calculated as normalized distance between two nodes, i.e. high weight connectivity were shown by low distance value. Core genes from STRING network graph was cross-observed in present studied graph (Table 1). Matched experimentally evidenced genes were: AIP, BTK, GNAS, GPR101, HESX1, LHCGR, LHX3, LHX4, MC2R, OTX2, PAX8, PDE4D, PRKAR1A, PROP1, SOX3, STAT5B, TBX19, and TSHR. While unmatched experimentally evidenced genes were: GHR, POU1F1. Comparative results from STRING network graph and Deep-learning based network graph (Table 2). It includes experimentally validated matching genes and possible genes which has highest possibilities to be experimentally validated in coming future. Possibilities for future experiments validation can be achieved for: ACTHR, AGMX1, ATK, BPK, DPDE3, ERBA2, FSHB, GH1, GHSR, GNAS1, GSP, HANF, LCGR, LGR2, LGR3, LHRHR, NR1A2, PKR1, PRKAR1, RNF216, SBP2, SECISBP2, THR1, THRB, TPIT, TRIAD3, TSE1, UBCE7IP1, XAP2, and ZIN. Separate network clusters: HIGHLY linked and LOW linked were shown in Figure 4 (a to l).

**Figure 2.**
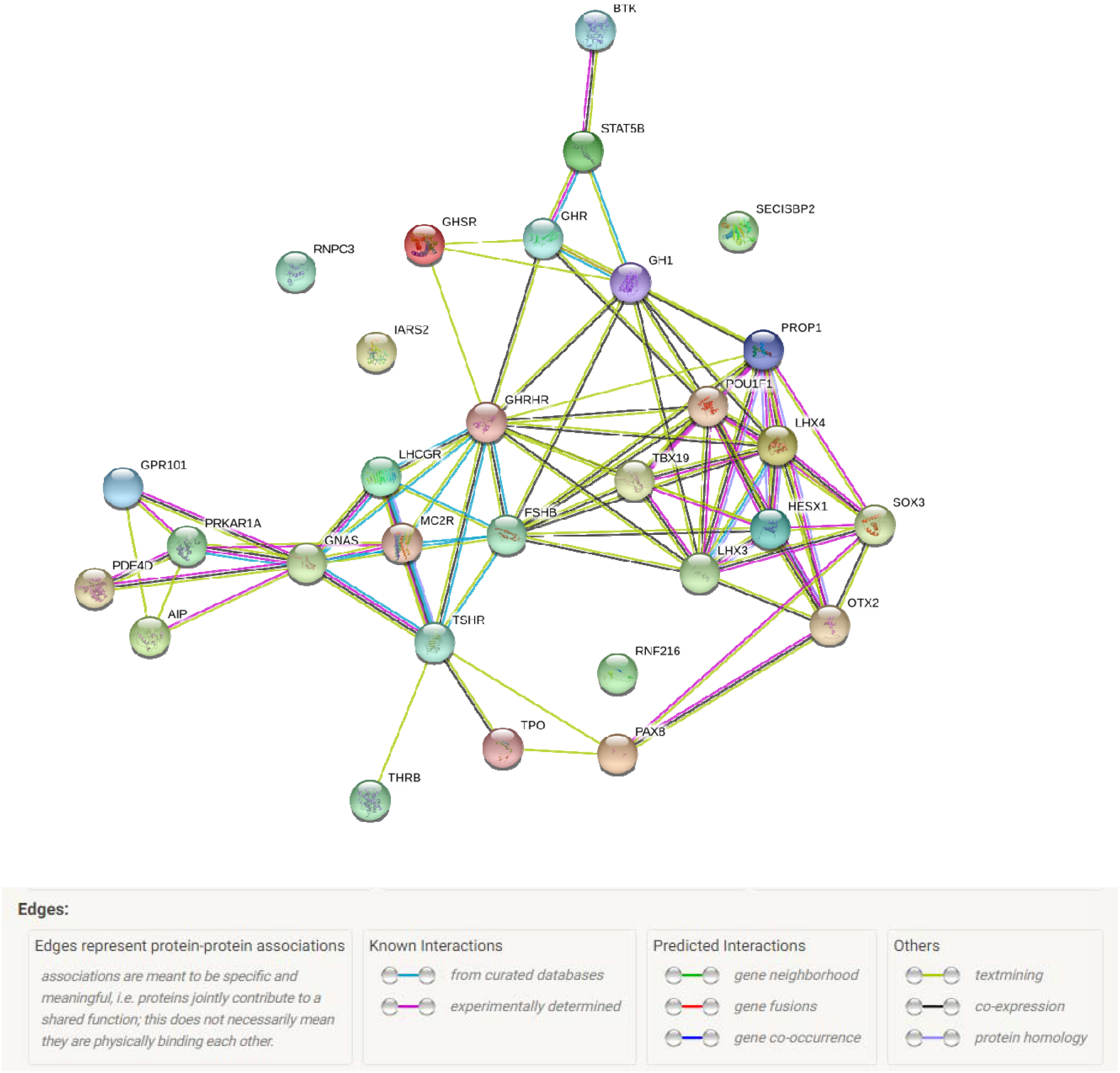
Reference network (from STRING database) for cross validation of developed protocol

**Figure 3.**
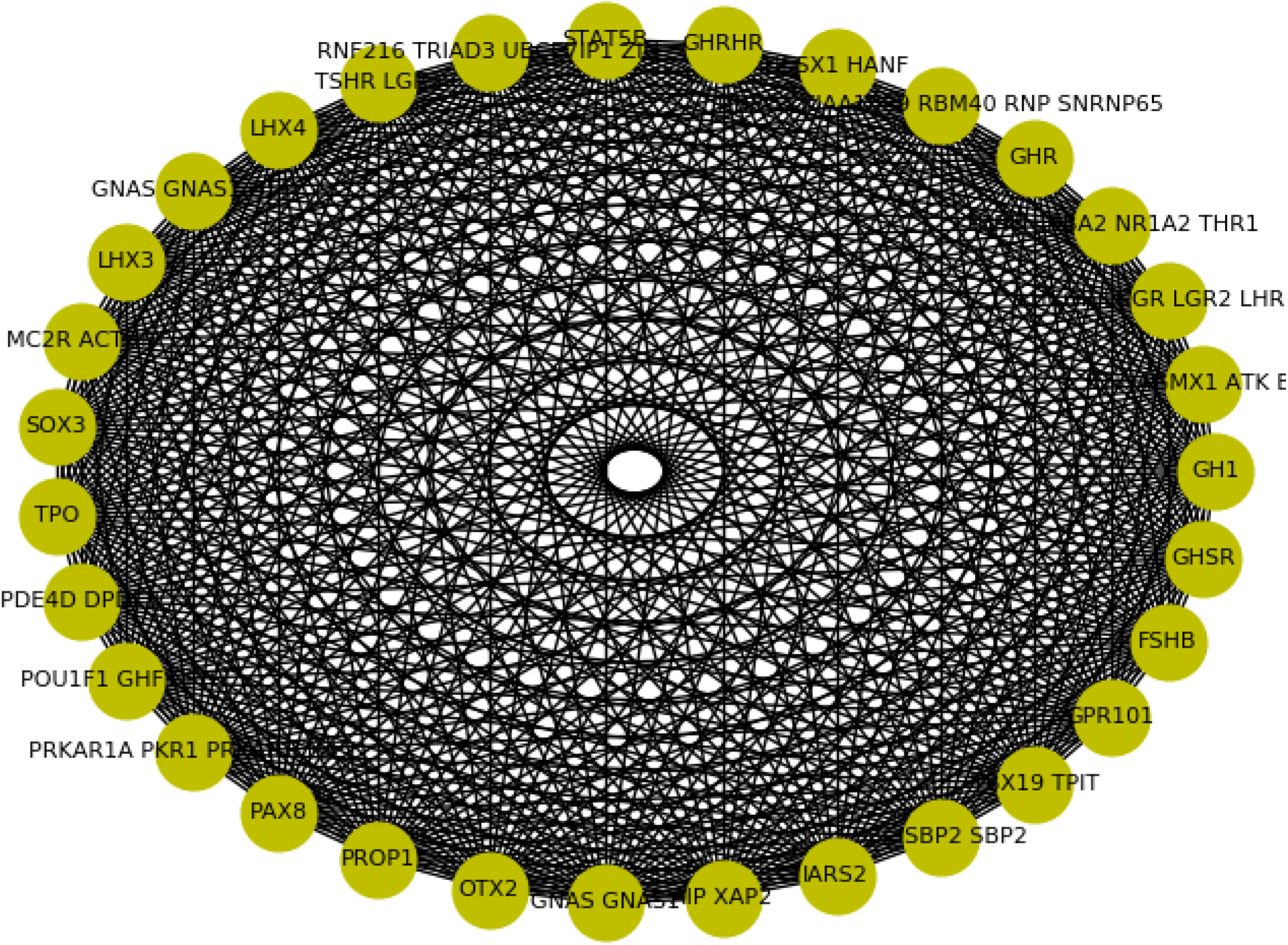
Network circular graph with all combinations including edge weight of 0 to 1; Here Node value was calculated through deep-learning based multiple models; Inter-node edge weight was calculated as normalized distance between two nodes, i.e. high weight connectivity were shown by low distance value.

**Figure 4(a).**
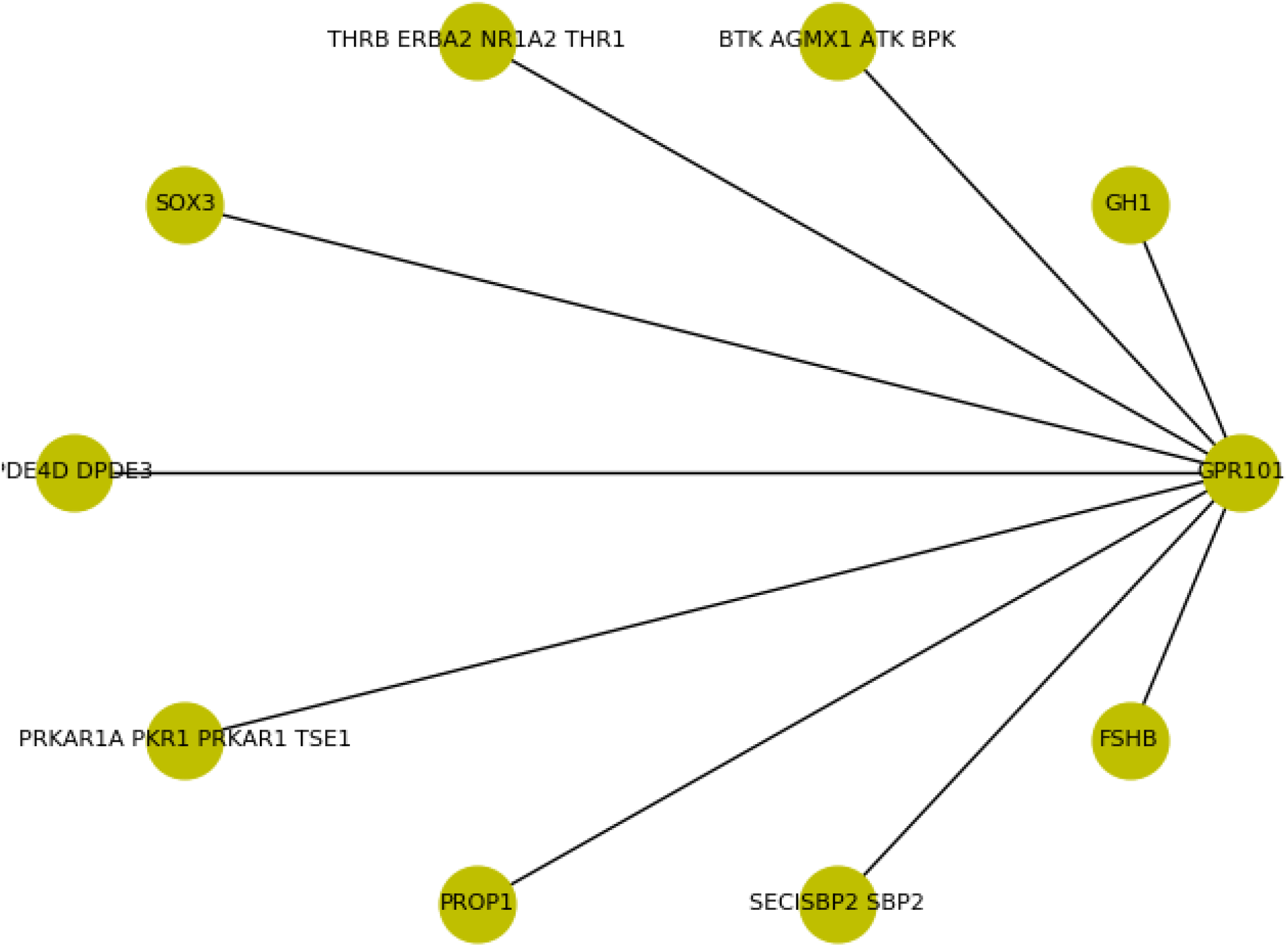
GPR101 centered cluster with in node distance < 0.05. It has highly evident clustered genes.

**Figure 4(b).**
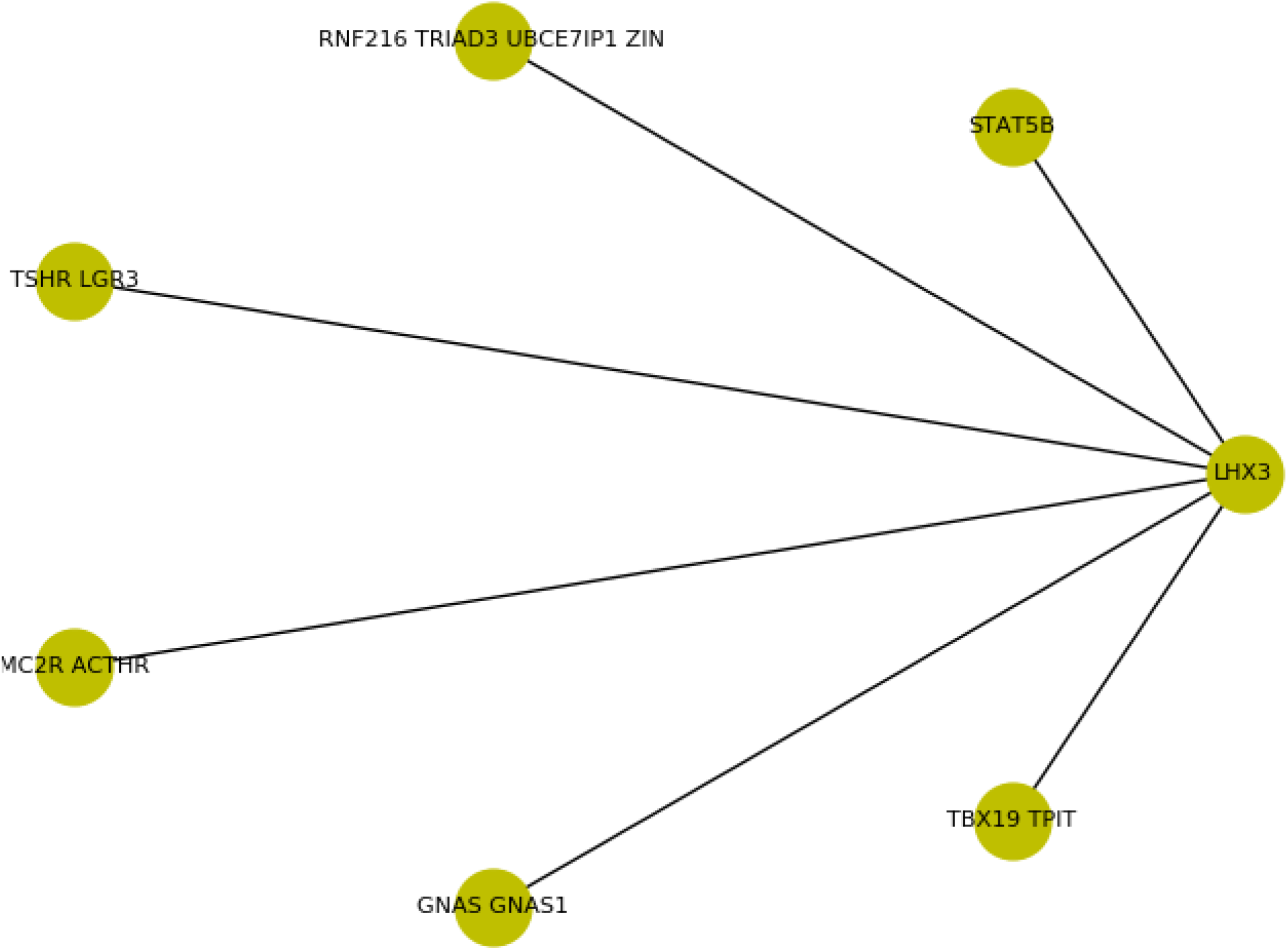
LHX3 centered cluster with in node distance < 0.05. It has highly evident clustered genes.

**Figure 4(c).**
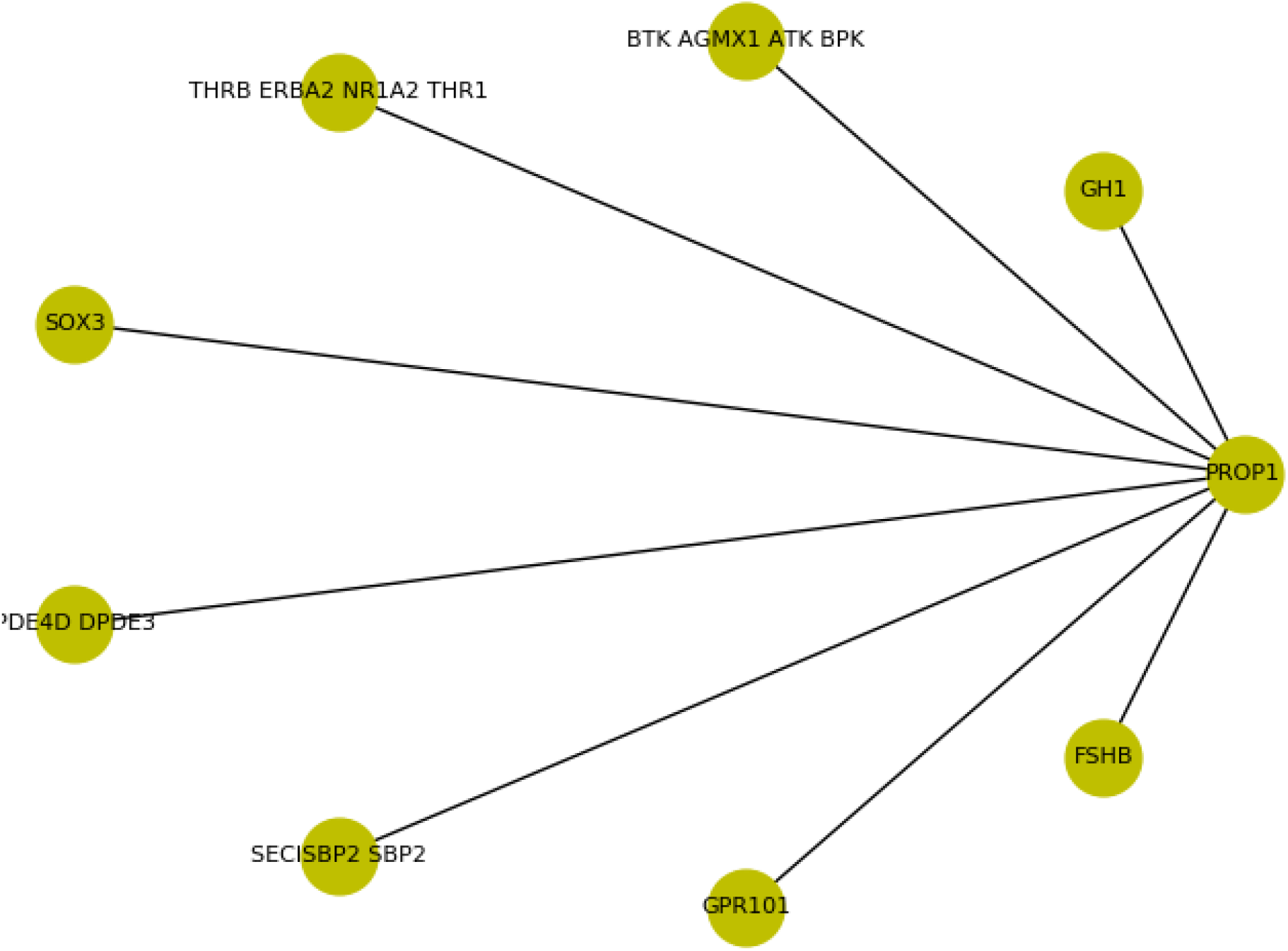
PROP1 centered cluster with in node distance < 0.05. It has highly evident clustered genes.

**Figure 4(d).**
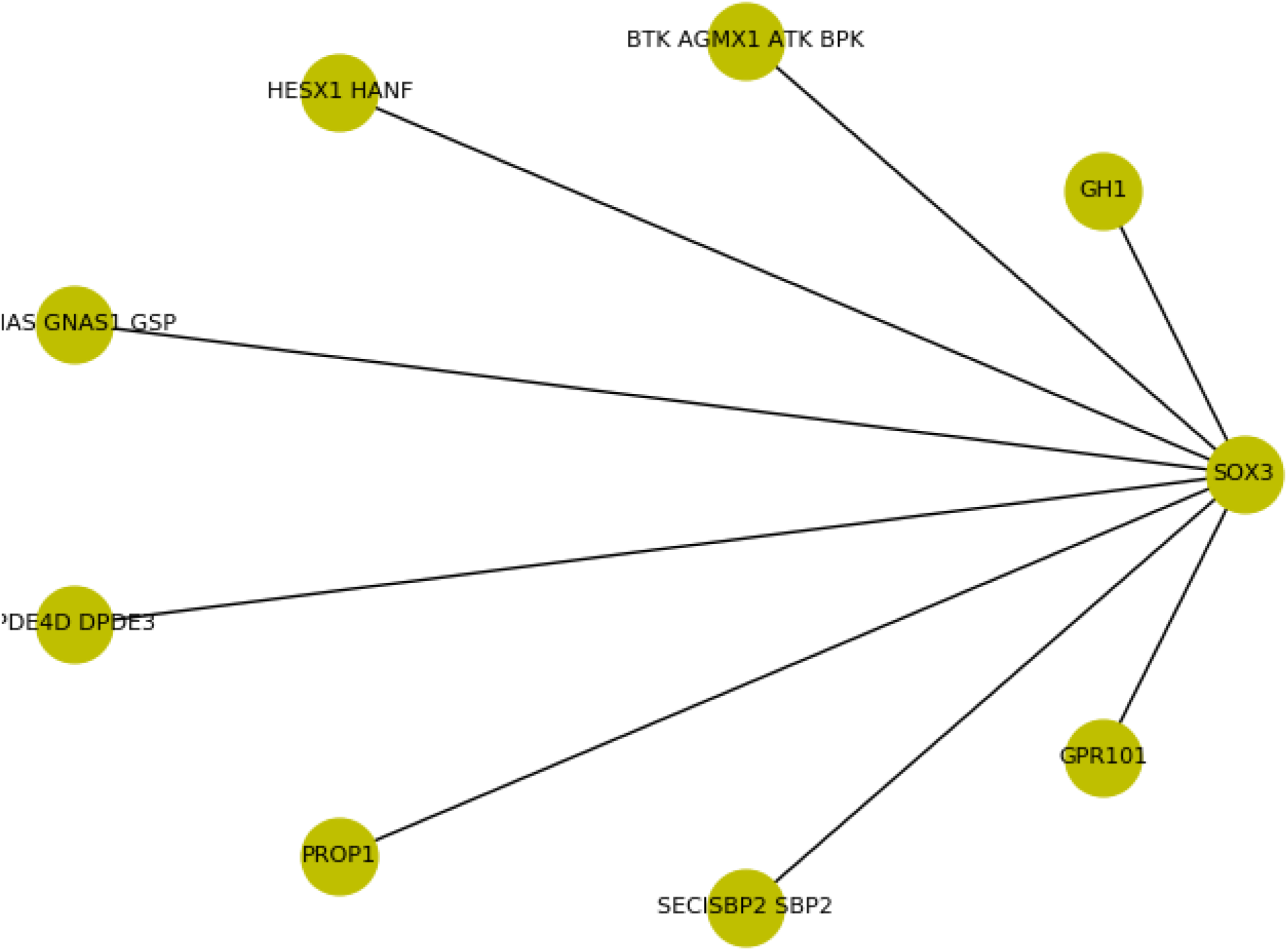
SOX3 centered cluster with in node distance < 0.05. It has highly evident clustered genes.

**Figure 4(e).**
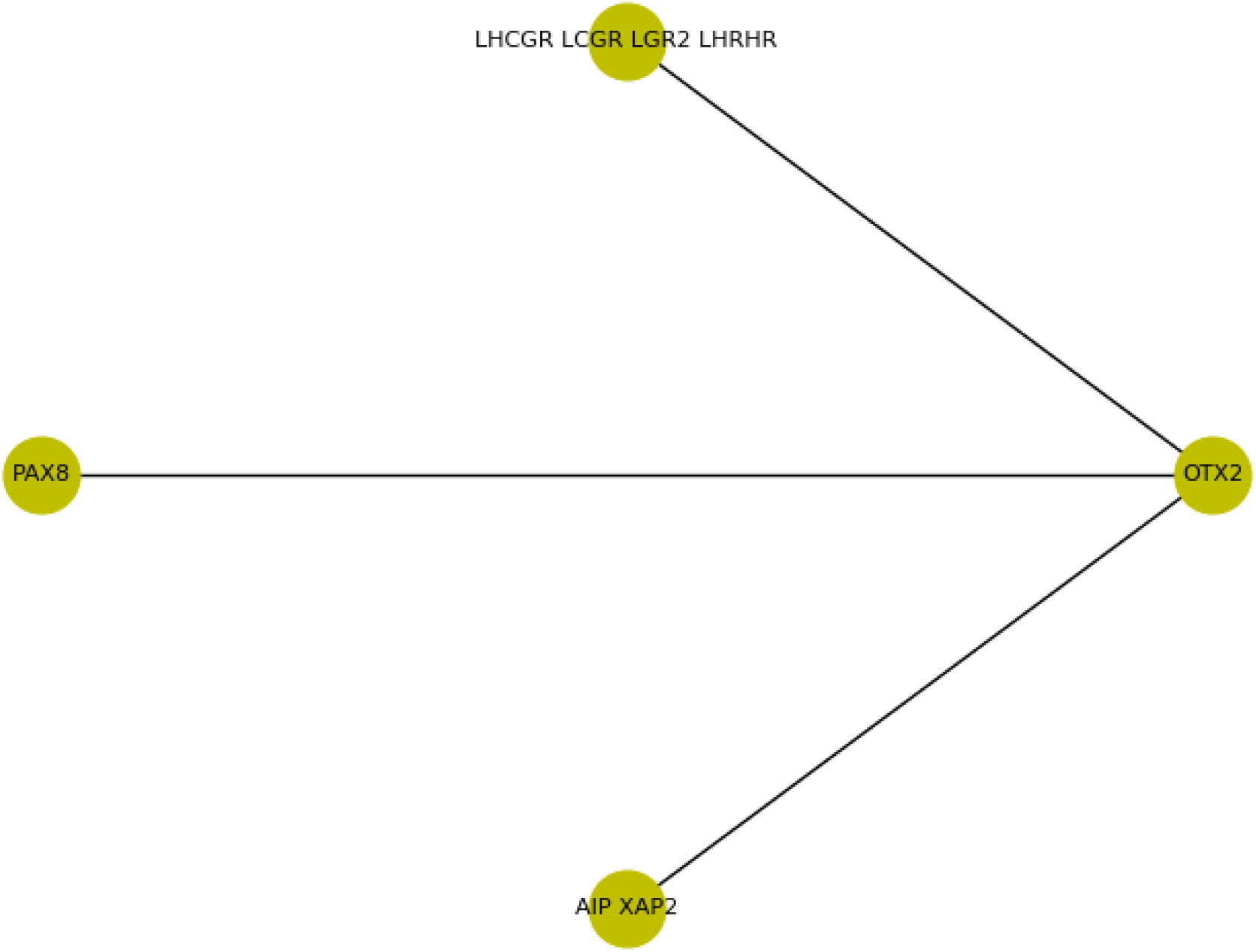
OTX2 centered cluster with in node distance < 0.05. It has highly evident clustered genes.

**Figure 4(f).**
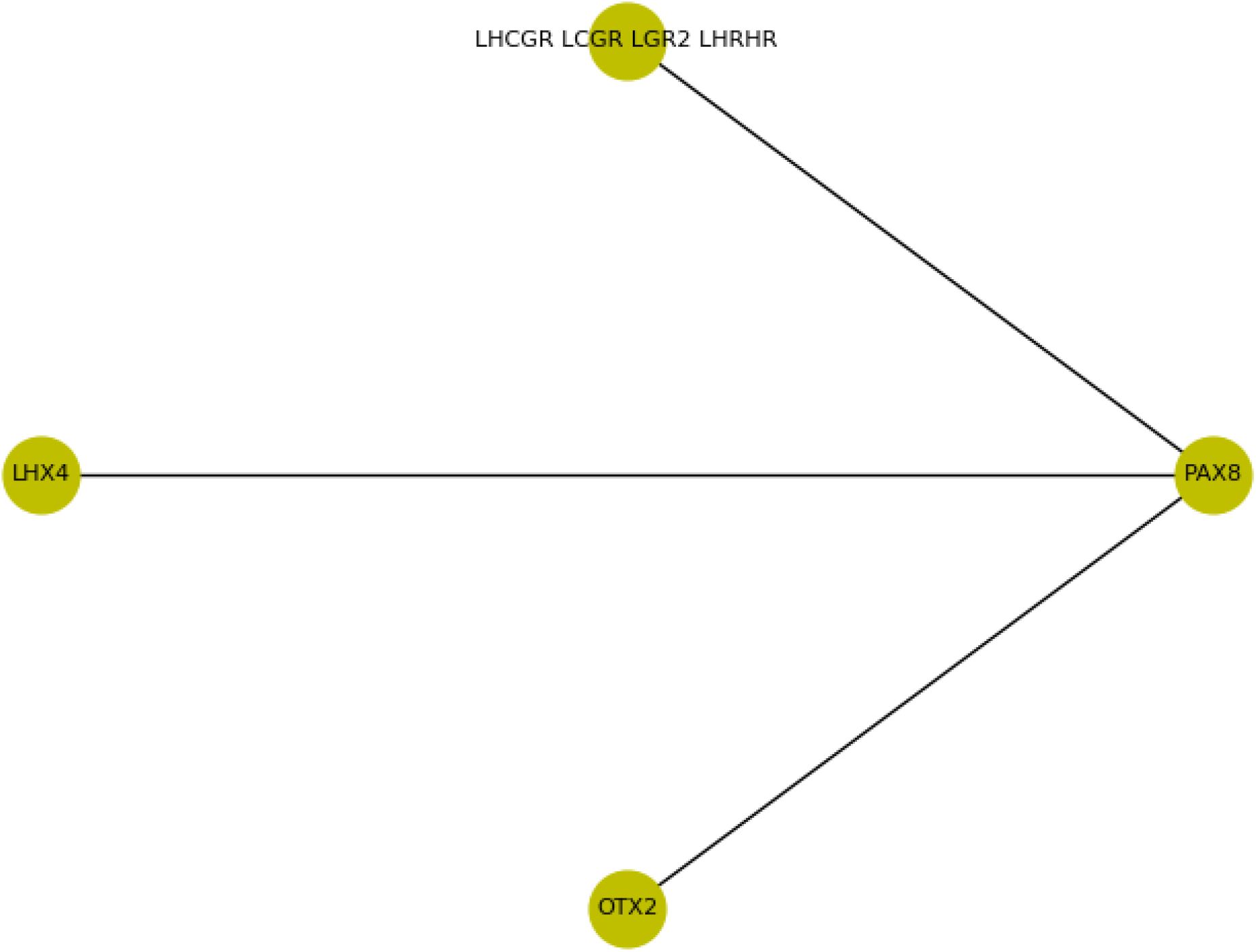
PAX8 centered cluster with in node distance < 0.05. It has highly evident clustered genes.

**Figure 4(g).**
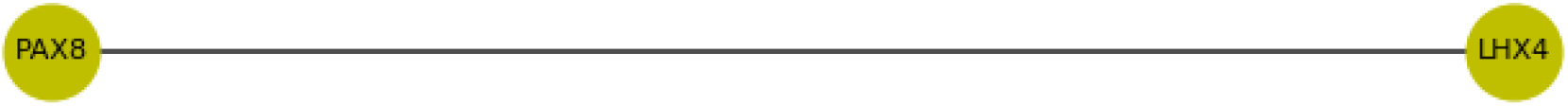
LHX4 centered cluster with in node distance < 0.05. It has highly evident clustered genes.

**Figure 4(h).**
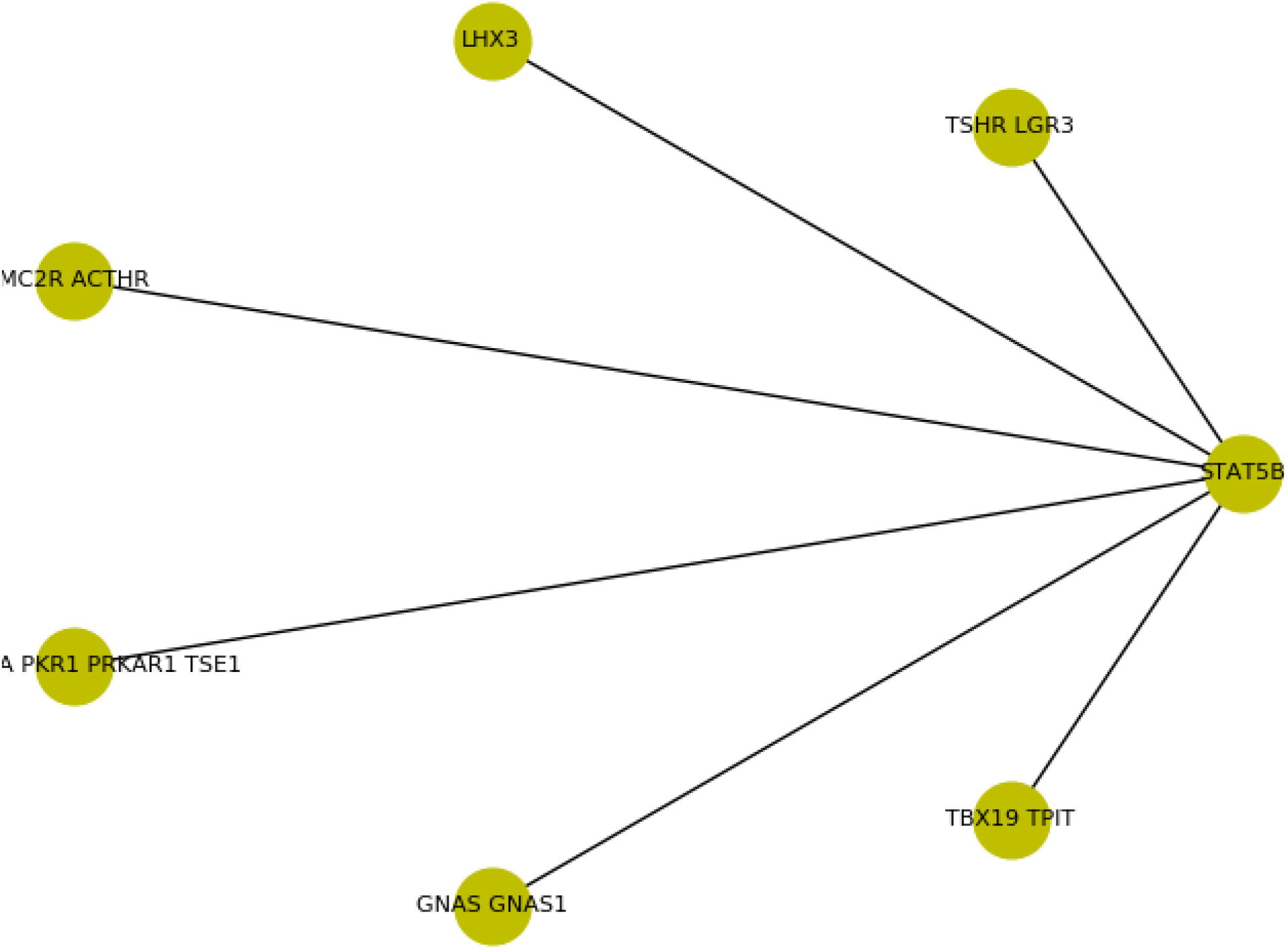
STAT5B centered cluster with in node distance < 0.05. It has highly evident clustered genes.

**Figure 4(i).**
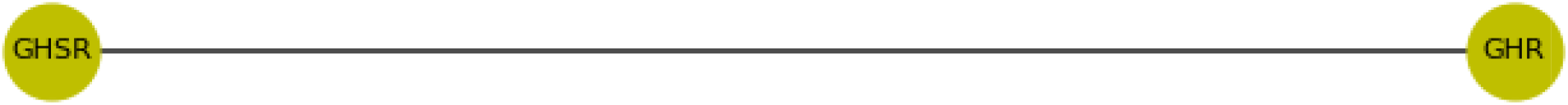
GHR centered cluster with in node distance < 0.05. It has highly evident clustered genes.

**Figure 4(j).**
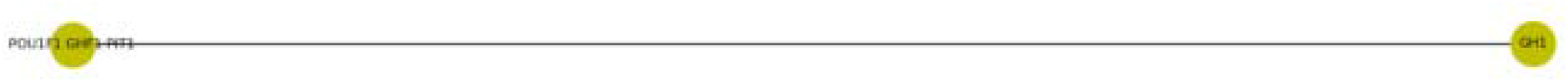
GH1 centered cluster with in node distance > 0.8. It has LOW evident clustered genes.

**Figure 4(k).**
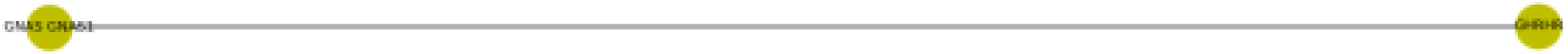
GHRHR centered cluster with in node distance of 0.8. It has LOW evident clustered genes.

**Figure 4(l).**
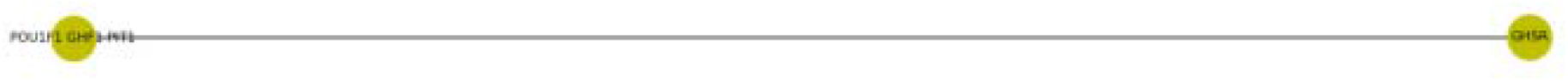
GHR centered cluster with in node distance > 0.9. It has LOW evident clustered genes.

**Table 1.** Core genes from STRING network graph was cross-observed in present studied graph

| HIGH evidenced Genes | Present study | LOW evidenced Genes | Present study |
| --- | --- | --- | --- |
| Gene from STRING graph with at least 01 links of experimental evidences | Found in LOW distant range (i.e. HIGH possibility) | Gene from STRING graph with NO links of experimental evidences | Found in HIGH distant range (i.e. LOW possibility) |
| PROP1 | <0.05: 'GH1', 'BTK AGMX1 ATK BPK', 'THRB ERBA2 NR1A2 THR1', 'SOX3', 'PDE4D DPDE3', 'SECISBP2 SBP2', 'GPR101', 'FSHB' | GHSR | >0.90: 'POU1F1 GHF1 PIT1' |
| POU1F1 | No in 20% | GH1 | >0.80: 'POU1F1 GHF1 PIT1' |
| LHX4 | <0.05: PAX8 | SECISBP2 | No till 0.80 |
| TBX19 | No in 20% | IARS2 | No till 0.80 |
| HESX1 | No in 20% | GHRHR | >0.80: 'GNAS GNAS1' |
| LHX3 | <0.05: 'STAT5B', 'RNF216<br>TRIAD3 UBCE7IP1 ZIN', 'TSHR<br>LGR3', 'MC2R ACTHR', 'GNAS<br>GNAS1', 'TBX19 TPIT' | RNPC3 | No till 0.80 |
| SOX3 | <0.05: 'GH1', 'BTK AGMX1 ATK<br>BPK', 'HESX1 HANF', 'GNAS<br>GNAS1 GSP', 'PDE4D DPDE3',<br>'PROP1', 'SECISBP2 SBP2',<br>'GPR101' | FSHB | No till 0.80 |
| OTX2 | <0.05: 'LHCGR LCGR LGR2<br>LHRHR', 'PAX8', 'AIP XAP2' | THRB | No till 0.80 |
| PAX8 | <0.05: 'LHCGR LCGR LGR2<br>LHRHR', 'LHX4', 'OTX2' | TPO | No till 0.80 |
| GNAS | No in 20% | RNF216 | No till 0.80 |
| PRKAR1A | No in 20% |  |  |
| BTK | No in 20% |  |  |
| STAT5B | <0.05: 'TSHR LGR3', 'LHX3',<br>'MC2R ACTHR', 'PRKAR1A PKR1<br>PRKAR1 TSE1', 'GNAS GNAS1',<br>'TBX19 TPIT' |  |  |
| GHR | <0.05: GHSR |  |  |
| AIP | No in 20% |  |  |
| PDE4D | No in 20% |  |  |
| GPR101 | 'GH1', 'BTK AGMX1 ATK BPK',<br>'THRB ERBA2 NR1A2 THR1',<br>'SOX3', 'PDE4D DPDE3',<br>'PRKAR1A PKR1 PRKAR1 TSE1',<br>'PROP1', 'SECISBP2 SBP2', 'FSHB' |  |  |
| LHCGR | No in 20% |  |  |
| MC2R | No in 20% |  |  |
| TSHR | No in 20% |  |  |

**Table 2.**
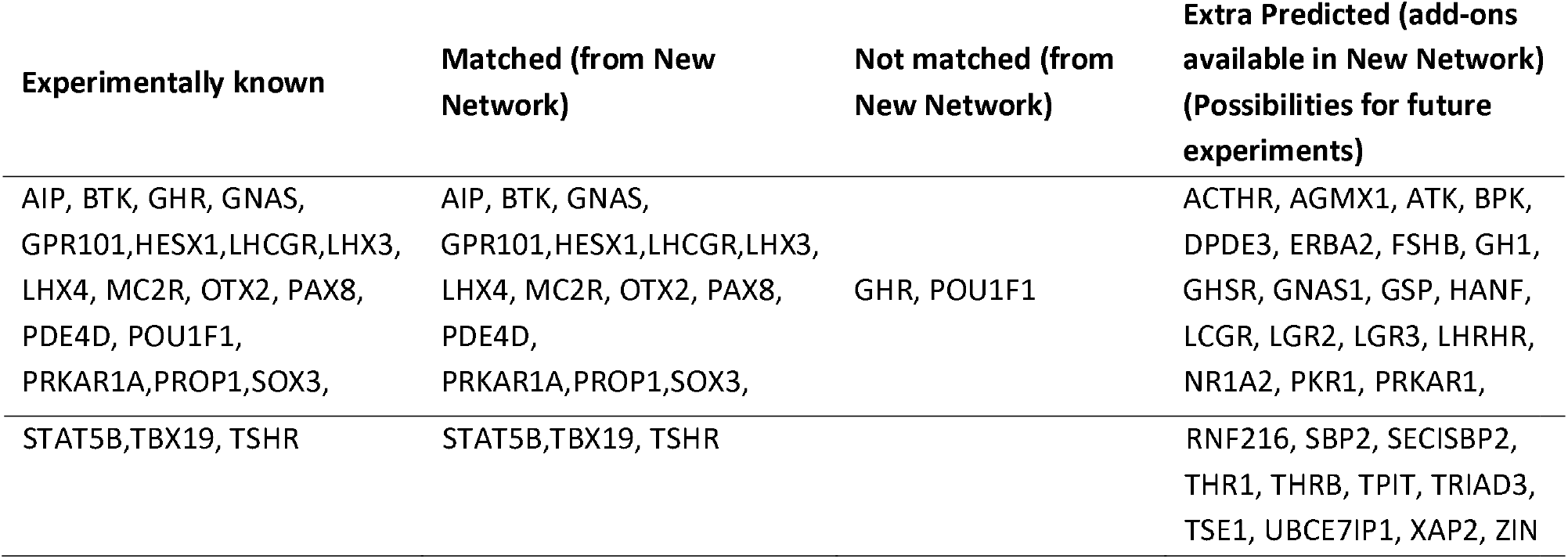
Comparative results from STRING network graph and Deep-learning based network graph

## CONCLUSION

Protocol established and validated for clustering of non-unified protein sequences through memory-map guided deep learning. Possibilities for future experiments validation have been achieved for genes named as: ACTHR, AGMX1, ATK, BPK, DPDE3, ERBA2, FSHB, GH1, GHSR, GNAS1, GSP, HANF, LCGR, LGR2, LGR3, LHRHR, NR1A2, PKR1, PRKAR1, RNF216, SBP2, SECISBP2, THR1, THRB, TPIT, TRIAD3, TSE1, UBCE7IP1, XAP2, and ZIN.

## ACKNOWLEDGEMENT

Author express gratitude to *The Institute of Mathematical Sciences*, Chennai-600113, India for providing research facilities as well as DAE Post-Doctoral Fellowship (PDF 214).

---

**Supplementary Table 1.**

| UniProt ID | Gene names | Protein names |
| --- | --- | --- |
| P01241 | GH1 | Somatotropin (Growth hormone) (GH) (GH-N) (Growth hormone 1) (Pituitary growth hormone) |
| Q06187 | BTK AGMX1 ATK BPK | Tyrosine-protein kinase BTK (EC 2.7.10.2) (Agammaglobulinemia tyrosine kinase) (ATK) (B-cell progenitor kinase) (BPK) (Bruton tyrosine kinase) |
| P22888 | LHCGR LCGR LGR2 LHRHR | Lutropin-choriogonadotropic hormone receptor (LH/CG-R) (Luteinizing hormone receptor) (LHR) (LSH-R) |
| P10828 | THRB ERBA2 NR1A2 THR1 | Thyroid hormone receptor beta (Nuclear receptor subfamily 1 group A member 2) (c-erbA-2) (c-erbA-beta) |
| P10912 | GHR | Growth hormone receptor (GH receptor) (Somatotropin receptor) [Cleaved into: Growth hormone-binding protein (GH-binding protein) (GHBP) (Serum-binding protein)] |
| Q96LT9 | RNPC3 KIAA1839 RBM40 RNP SNRNP65 | RNA-binding region-containing protein 3 (RNA-binding motif protein 40) (RNA-binding protein 40) (U11/U12 small nuclear ribonucleoprotein 65 kDa protein) (U11/U12 snRNP 65 kDa protein) (U11/U12-65K) |
| Q9UBX0 | HESX1 HANF | Homeobox expressed in ES cells 1 (Homeobox protein ANF) (hAnf) |
| Q02643 | GHRHR | Growth hormone-releasing hormone receptor (GHRH receptor) (Growth hormone-releasing factor receptor) (GRF receptor) (GRFR) |
| P51692 | STAT5B | Signal transducer and activator of transcription 5B |
| Q9NWF9 | RNF216 TRIAD3 UBCE7IP1 ZIN | E3 ubiquitin-protein ligase RNF216 (EC 2.3.2.27) (RING finger protein 216) (RING-type E3 ubiquitin transferase RNF216) (Triad domain-containing protein 3) (Ubiquitin-conjugating enzyme 7-interacting protein 1) (Zinc finger protein inhibiting NF-kappa-B) |
| P16473 | TSHR LGR3 | Thyrotropin receptor (Thyroid-stimulating hormone receptor) (TSH-R) |
| Q969G2 | LHX4 | LIM/homeobox protein Lhx4 (LIM homeobox protein 4) |
| P63092 | GNAS GNAS1 GSP | Guanine nucleotide-binding protein G(s) subunit alpha isoforms short (Adenylate cyclase-stimulating G alpha protein) |
| Q9UBR4 | LHX3 | LIM/homeobox protein Lhx3 (LIM homeobox protein 3) |
| Q01718 | MC2R ACTHR | Adrenocorticotrophic hormone receptor (ACTH receptor) (ACTH-R) (Adrenocorticotropin receptor) (Melanocortin receptor 2) (MC2-R) |
| P41225 | SOX3 | Transcription factor SOX-3 |
| P07202 | TPO | Thyroid peroxidase (TPO) (EC 1.11.1.8) |
| Q08499 | PDE4D DPDE3 | cAMP-specific 3',5'-cyclic phosphodiesterase 4D (EC 3.1.4.53) (DPDE3) (PDE43) |
| P28069 | POU1F1 GHF1 PIT1 | Pituitary-specific positive transcription factor 1 (PIT-1) (Growth hormone factor 1) (GHF-1) |
| P10644 | PRKAR1A PKR1 PRKAR1 TSE1 | cAMP-dependent protein kinase type I-alpha regulatory subunit (Tissue-specific extinguisher 1) (TSE1) |
| Q06710 | PAX8 | Paired box protein Pax-8 |
| O75360 | PROP1 | Homeobox protein prophet of Pit-1 (PROP-1) (Pituitary-specific homeodomain factor) |
| P32243 | OTX2 | Homeobox protein OTX2 (Orthodenticle homolog 2) |
| P84996 | GNAS GNAS1 | Protein ALEX (Alternative gene product encoded by XL-exon) |
| O95467 | GNAS GNAS1 | Neuroendocrine secretory protein 55 (NESP55) [Cleaved into: LHAL tetrapeptide; GPIPIRRH peptide] |
| O00170 | AIP XAP2 | AH receptor-interacting protein (AIP) (Aryl-hydrocarbon receptor-interacting protein) (HBV X-associated protein 2) (XAP-2) (Immunophilin homolog ARA9) |
| Q9NSE4 | IARS2 | Isoleucine--tRNA ligase, mitochondrial (EC 6.1.1.5) (Isoleucyl-tRNA synthetase) (IleRS) |
| Q96T21 | SECISBP2 SBP2 | Selenocysteine insertion sequence-binding protein 2 (SECIS-binding protein 2) |
| O60806 | TBX19 TPIT | T-box transcription factor TBX19 (T-box protein 19) (T-box factor, pituitary) |
| Q96P66 | GPR101 | Probable G-protein coupled receptor 101 |
| Q5JWF2 | GNAS GNAS1 | Guanine nucleotide-binding protein G(s) subunit alpha isoforms XLas (Adenylate cyclase-stimulating G alpha protein) (Extra large alphas protein) (XLalphas) |
| P01225 | FSHB | Follitropin subunit beta (Follicle-stimulating hormone beta subunit) (FSH-B) (FSH-beta) (Follitropin beta chain) |
| Q92847 | GHSR | Growth hormone secretagogue receptor type 1 (GHS-R) (GH-releasing peptide receptor) (GHRP) (Ghrelin receptor) |

